# Discovery of microbial glycoside hydrolases via enrichment and metaproteomics

**DOI:** 10.1101/2025.02.11.637619

**Authors:** Jitske M. van Ede, Suzanne van der Steen, Geert M. van der Kraan, Mark C.M. van Loosdrecht, Martin Pabst

## Abstract

The immense microbial diversity on Earth represents a vast genomic resource, yet discovering novel enzymes from complex environments remains challenging. Here, we combine microbial enrichment with metagenomics and metaproteomics to facilitate the identification of microbial glycoside hydrolases that operate under defined conditions. We enriched microbial communities using the carbohydrate polymer pullulan at elevated temperatures and acidic conditions. Pullulan is a natural polysaccharide composed of maltotriose units linked by α-1,6 glycosidic bonds. Along with its hydrolyzing enzymes, it has broad applications across various industries. The enrichment inocula were sampled from thermophilic compost and soil from the bank of a pond. In both cases, Alicyclobacillus emerged as the dominant microorganism. Metaproteomic analysis of the enrichment biomass and secretome identified several pullulan-degrading enzymes from this organism. Notably, these enzymes were absent in the metagenomic analysis of the initial inoculum, underscoring the effectiveness of combining microbial enrichment with multi-omics for uncovering novel enzymes from complex microbial environments.

## Introduction

Enzymes are natural biocatalysts, which have the potential to significantly contribute to sustainable processes in biotechnology and industry. Not only are they biodegradable, enzymes also have the potential to conduct conversions under ambient conditions, reduce the use of fossil fuels and chemical waste. Moreover, enzymes provide complex molecules that are challenging to produce via chemical synthesis alone^1,2^. Enzymes are already extensively employed in industry, with hydrolases being the most prevalent class of enzymes in use^3^. Hydrolases can modify biopolymers to alter their properties or break them down into smaller fragments, with a vast array of applications existing for (modified) biopolymers. For example, carbohydrate biopolymers are widely applied in the food industry, nutraceuticals, and care products. Cellulose, obtained from plant cell walls, is used as a stabilizer and thickener in various food formulations^4^. Chitosan, derived from crustacean shells, exhibits antimicrobial properties, making it a valuable component in dietary supplements^5^. Pullulan, a natural polysaccharide produced by the fungus *Aureobasidium pullulans*, is highly valued in the food and pharmaceutical industries for its ability to form edible, non-toxic films and coatings^6,7^. Its applications include encapsulating food flavours and fragrances, improving product shelf life, and acting as a binder in orally dissolvable films for pharmaceuticals, demonstrating its versatility and biocompatibility^8^. Furthermore, also the hydrolysing enzymes are used in industrial applications. For example, pullulanases are essential in the starch-processing industry, where they are used to hydrolyze pullulan into maltotriose units, thereby enhancing the efficiency of starch liquefaction and saccharification processes. Their use significantly improves the production of glucose and maltose syrups, providing a higher yield and specificity compared to conventional methods^9^. Hydrolases are also employed in cleaning processes, such as laundry. These enzymes facilitate the breakdown of polymers, thereby boosting cleaning efficiency. Additionally, their application offers environmental benefits by enabling lower washing temperatures, shorter wash times, and reduced water usage^10,11^.

Pullulan represents a polysaccharide which consists of maltotriose units, in which three glucoses are linked via an alpha-1,4 linkage, and these maltotriose units are linked via alpha-1,6 glycosidic bonds^12^. Pullulan degrading enzymes can be divided into two main groups, pullulanases (type I and type II) and pullulan hydrolases (type I, II and III). Pullulanases cleave the α-1,6 bond, thereby converting pullulan into maltotriose. In addition, pullulanases are also active against other sugar polymers, including starch and glycogen. While pullulanase type I can only cleave the α-1,6 linkages of these sugar polymers, pullulanase type II can cleave both the α-1,6 and α-1,4 linkages. Pullulan hydrolases are further divided in three types. Type I (neopullulanase) and type II (isopullulanase) cleave the α-1,4 linkage of pullulan, thereby forming panose and isopanose, respectively. Type III has the ability to cleave both the α-1,4 and α-1,6 linkage of pullulan, forming panose, maltotriose, maltose and glucose^13^. Although a wide variety of enzymes are utilized in various industries, the quest for new enzymes–including pullulanases–with improved features is ongoing^13^. There is currently a focused effort on employing advanced protein/enzyme optimization techniques, such as artificial intelligence, directed evolution, and intelligent design, to improve enzymes^14-20^. However, despite progress in protein engineering and other modification strategies, these approaches don’t always yield the anticipated results due to the complex relationship between the structure and function of proteins^21^. Fortunately, a remarkable diversity of microorganisms inhabit our planet, providing us with a vast genetic, and therefore biocatalytic resource^2^. Exploring this natural diversity with sophisticated enzyme discovery methods presents a promising alternative for finding enzymes with the desired traits or for identifying templates for further engineering.

Conservative methods to mine genetic resources use cultivation of a microorganism and to clean streak it on a plate under certain conditions. Subsequently, a colony is isolated of which the DNA is extracted and sequenced^22,23^. After the genome is assembled it can be mined for genetic targets of interest. Nevertheless, this approach has limitations, namely, most microorganisms cannot be grown as a pure culture under conditions which prevail in laboratory environments. This limitation is better known as “the great plate anomaly”. It is expected that only about 1% of the microorganisms in a sampled environment can be found in a lab after enrichment^24-26^. A way to avoid this problem and become culture independent is by using metagenomics^27^. Over the past years, whole metagenome sequencing became a popular method to sequence the entire metagenome from natural microbial communities^28^. Currently, two main metagenomic strategies can be distinguished, the sequence based and the function based approach^29^. The function-based approach involves constructing a genomic library from environmental DNA, followed by screening individual genes in a cultivable host cell for desired activities^28-30^. While this method enables the discovery of entirely novel enzymes, it is limited by high costs and the requirement of high-throughput activity assays^2^. In contrast, the sequence based approach depends on sequence similarity to known templates^29,30^, which restricts its ability to identify enzymes with novel functions^2^.

More recent strategies include microbial enrichment cultures followed by multi-omics analysis of the cultures (genomics and proteomics, supported by advanced bioinformatics tools)^31^. These strategies first enrich for certain organisms which possess an enzyme with a desired function. This route is only feasible when the enzyme plays a crucial role in the growth of the organism, e.g. is involved in the carbon or nitrogen source utilization^32^. For instance, in pursuit of hydrolytic enzymes which cleave specific polymers, an enrichment with the polymer as the sole carbon source can be performed. This ensures that only microorganisms that can hydrolyse and utilize the polymer hydrolysis products will grow^33^. Moreover, the conditions for enriching microbes can be tuned towards the conditions in which the enzymes should be active (e.g. temperature, pH and salinity). Subsequently, instead of only employing metagenomics to mine for the enzymes of interest, a more streamlined approach is to also integrate proteomics into the workflow^31,34^. Advantageously, metaproteomics allows to identify the actually expressed enzymes, as well as their cellular location, such as cytosolic, membrane-bound, or secreted^35,36^. This greatly simplifies genomic data and it supports identification of novel enzyme candidates or those with enhanced properties. It also provides template sequences for further engineering to optimize desired properties^37^. In summary, this study demonstrates the discovery of novel pullulan-degrading enzymes that operate at defined conditions, by combining microbial enrichment with metagenomics and metaproteomics.

## Methods

### Microbial enrichment

Organic matter from compost was obtained from a compost pile in Waddinxveen (further referred to as “compost”), The Netherlands. Soil was collected from two locations in Delft, The Netherlands (at approximately 51°59’36.3”N 4°22’23.6”E and 51°59’37.5”N 4°22’34.0”E) and combined (further referred to as “soil”). The enrichments were performed under sterile conditions (to ensure only microbial diversity from the sampled sources are enriched) in 250 ml shake flasks with screw caps. The growth medium contained 0.5 g/L KH_2_PO_4_, 3 g/L (NH_4_)_2_SO_4_, 0.2 g/L MgSO_4_•7H_2_O, 10 ml/L MD-VS ATCC vitamin supplement and 2.5 ml/L trace element solution (15 g/L EDTA (Titriplex III), 4.5 g/L ZnSO_4_•7H_2_O, 0.84 g/L MnCl_2_•2H_2_O, 0.30 CoCl_2_•6H_2_O, 0.20 g/L CuSO_4_•5H_2_O, 0.40 g/L Na-_2_MoO_4_•2H_2_O, 4.5 g/L CaCl_2_•2H_2_O, 3.0 g/L FeSO_4_•7H_2_O, 1.0 g/L H_3_BO_3_ and 0.10 g/L KI). Pullulan was added to a final concentration of 10 g/L and the pH of the medium was adjusted to 4.5 with 2 M HCl. Finally, the medium was sterilized in the autoclave at 110 °C for 30 min. and subsequently stored at 4 °C until usage. 50 ml of growth medium was transferred to a 250 ml sterile flask. The medium was inoculated by either 0.5 g of compost further referred to as “compost enrichment”, or by 0.5 g of soil, further referred to as “soil enrichment”. In addition, a control culture without inoculum was included. The enrichments were kept at 57 °C, 100 rpm in the dark in an incubator hood (TH 15, Edmund Buhler GmbH). The pH was monitored twice a day using a pH probe (826 pH mobile, Metrohm) and if necessary adjusted back to pH 4.5 using 0.2 M KOH or 0.2 M HCl.

No organic buffer was used to prevent it from becoming a potential substrate for microbial growth. All steps were performed next to a Bunsen burner to maintain sterile conditions. The enrichments were transferred twice in a 1:1000 (v/v) ratio to fresh medium when almost all pullulan was consumed, after 3 and 4 days, respectively. After the second transfer, the OD_600_ (using the cell density meter Ultrospec 10, Biochrom) and the pullulan consumption (measured by HPLC analysis) were monitored daily, while the pH was still measured twice a day. The microbial enrichments were analyzed once all pullulan was consumed, after 8 and 14 days for the compost and soil enrichment, respectively. Images of the enrichments were created using the Zeiss Axio microscope with a 100x magnification.

In addition, a 250 ml flask was inoculated with 0.25 g compost combined with 0.25 g soil collected on the TUD campus and incubated at 75 °C in a water bath. The enrichment was monitored as described above. **Monitoring degradation of pullulan by HPLC**. Approximately 1 mL of culture broth was centrifuged for 10 min. at 4 °C, 14000 rcf, after which 667 μL of the supernatant was transferred to a clean screw cap vial. Trifluoroacetic acid (TFA) was added to reach a final concentration of 4 M TFA. The samples were incubated for 4 hours at 100 °C in the ThermoMixer with Thermotop (Eppendorf, Thermomixer C). After a centrifugation step of 15 min. at 4 °C, 14000 rcf, the supernatant was collected in a clean Eppendorf tube. The samples were measured on a Vanquish HPLC system (Thermo Scientific, Germany) with a Aminex HPX-87H separation column. A constant flow rate of 0.750 ml/min was maintained over a total run time of 45 minutes, using 1.5 mM phosphoric acid in milli-Q water as eluent. The column chamber temperature was kept at 50 °C and compounds were detected using the RI detector (ERC, RefractoMax520). Data analysis was performed with Chromeleon 7 (Thermo Scientific, Germany). **Whole metagenome sequencing**. Whole metagenome sequencing of the starting biomass and enrichments was performed by BaseClear B.V. (Leiden, The Netherlands). Briefly, DNA was extracted using standard molecular biology kits from Zymo Research, and DNA concentration was confirmed using a nanodrop spectrophotometer. Sequence libraries were generated using Nextera XT DNA Library Preparation Kit. Paired-end sequence reads were generated using the Illumina NovaSeq PE150 system. The sequences generated were performed under accreditation according to the scope of BaseClear B.V. (L457; NEN-EN-ISO/IEC 17025). FASTQ read sequence files were generated using bcl2fastq version 2.20 (Illumina). Initial quality assessment was based on data passing the Illumina Chastity filtering. Subsequently, reads containing PhiX control signal were removed using an in-house filtering protocol. In addition, reads containing (partial) adapters were clipped (up to a minimum read length of 50 bp). The second quality assessment was based on the remaining reads using the FASTQC quality control tool version 0.11.8. Metagenome assembly has been performed using MEGAHIT 1.2.9. Contigs smaller than 1000 base pairs have been removed from final assembly. PROKKA was used to locate open reading frames (ORFs) on contigs, and to translate the ORFs to protein sequences. These sequences were used as reference database for the metaproteomic analysis. Taxonomic classification of ORFs was performed using the DIAMOND sequence aligner and the NCBI sequence reference database. A consensus linage for every ORF was obtained by using the bit score approach from the top 25 alignments described earlier^35,38^. Functional classification was performed as described below. In addition, a sequence alignment of the *CdaA* (Uniprot: Q9WX32) and *AmyA* (Uniprot: Q06307) gene of *Alicyclobacillus acidocaldarius* was performed against the metagenomics data of both the enrichment cultures and the inocula using the DIAMOND sequence aligner. **Metaproteomics** Biomass of the enrichment cultures were collected in a clean LoBind Eppendorf tube (approx. 25 mg wet weight). Subsequently, 0.175 mL of TEAB resuspension buffer (50 mM TEAB, 1% (w/w) sodium deoxycholate, pH 8), 0.175 mL of B-PER reagent (Bacterial Protein Extraction Reagent, Thermo Scientific) and 0.15 g of acid washed glass beads (150-212 μm) were added. The samples were homogenized using the bead beater for 1.5 min., followed by a 1-minute incubation on ice and a centrifugation for 2 min. at 4 °C, 14000 rcf. This cycle was repeated twice. Afterwards, the samples were subjected to an ultrasonic bath for 10 min. and centrifuged for 15 min. at 4 °C, 14000 rcf. All centrifugation steps were conducted at 4 °C and 14000 rcf unless stated otherwise. The supernatant was collected in a LoBind Eppendorf tube. Trichloroacetic acid (TCA) was added in a 1:4 ratio to the supernatant. The samples were vortexed and incubated for 30 min. at 4 °C, followed by centrifugation for 15 min. The supernatant was discarded, and the protein pellet was washed with 200 μL of ice-cold acetone, vortexed and centrifuged for 15 min. The supernatant was removed and 100 μL of 6 M Urea was added to the protein pellets. To re-dissolve the pellets, the samples were vortexed thoroughly and incubated at 37 °C, 300 rpm in the ThermoMixer with Thermotop (Eppendorf, Thermomixer). If necessary, an additional 100 μL of 6 M Urea was added to dissolve the pellets. The samples were reduced by incubating at 37 °C for 60 min. after adding 30 μL 10 mM dithiothreitol, and subsequently alkylated for 30 minutes, at RT in the dark, by adding 30 μL of 20 mM iodoacetamide. The samples were diluted with 200 mM ammonium bicarbonate to achieve a Urea concentration < 1 M. To 100 μL of the final sample volume, 5 μL of 0.1 μg/μL trypsin solution was added. After gently shaking, the samples were incubated at 37 °C overnight. The samples were cleaned and concentrated using the Oasis HLB 96-well μElution Plate with 2 mg sorbent per well, 30 μm (Waters, UK). In short, the columns were conditioned with 750 μL MeOH, equilibrated with 2x 500 μL MS-H_2_O and then loaded with the samples. Two washing steps with 350 μL 5% MeOH in MS-H_2_O were performed, followed by elution with 200 μL 2% formic acid in 80% MeOH and 200 μL 1 mM ammonium bicarbonate (ABC) in 80% MeOH. The eluates were collected in a LoBind Eppendorf tube. The samples were dried using the SpeedVac concentrator (Thermo Scientific) at 50 °C and stored at -20 °C until analysis by LC-MS. For LC-MS analysis, the dried samples were dissolved in 15 μL 3% ACN plus 0.1% trifluoroacetic acid (TFA) in MS-H_2_O, incubated for 30 min. at RT, and vortexed thoroughly. The samples were diluted to a final protein concentration of approximately 0.5 mg/ml, estimated at 280 nm using the NanoDrop ND-1000 spectrophotometer (Thermo Scientific), before proteomic analysis. Extracellular proteins from the secretome of the enrichment cultures were prepared as described in the following. Samples from the enrichment culture were centrifuged for 15 min. and the supernatant was collected. Several protein precipitation methods were employed including TCA precipitation as described above, an acetone precipitation, an acetone/salt precipitation and a filter assisted sample preparation (FASP). The latter three are described below. For the acetone precipitation 250 μL sample was mixed with 1250 μL ice-cold acetone, vortexed and stored at -20 °C for 30 min. The samples were centrifuged for 15 min. and the supernatant was removed. For the acetone/salt precipitation 50 μL of 3 M NaCl solution was added to 250 μL sample. Subsequently, 1200 μL acetone (RT) was added and the samples were mixed gently. The samples were incubated for 30 min. at RT and centrifuged for 15 min. The supernatant was carefully removed, and the protein pellet was washed with 400 μL acetone (RT). Another 15-minute centrifuging was performed, and the supernatant was removed. After protein precipitation, the protein pellet was re-dissolved in 100 μL of 6 M Urea and processed as described above. For the filter assisted sample preparation 200 μL 6 M Urea was added to a 10 kDa Microcon filter (Merk-Millipore), followed by a centrifugation of 30 min. at 14000 rcf and 20 °C. After discarding the flow-through, this step was repeated once. Unless stated otherwise, all centrifugations during FASP were conducted at 14000 rcf and 20 °C. Either 250 μL sample or 4 x 500 μL sample was loaded onto the filter followed by centrifuging for 30 min. The samples were then reduced by adding 30 μL of 10 mM DTT and 70 μL of 200 mM ammonium bicarbonate. After vortexing and incubating at 37 °C for 60 min., 30 μL of 20 mM IAA was added to alkylate the samples. Following a 30 min. incubation in the dark at RT, the samples were centrifuged for 30 min. 100 μL of 6 M Urea was added and centrifuged for 30 min. Subsequently, 100 μL of 200 mM ammonium bicarbonate was added and centrifuged for 30 min.; this step was repeated twice. The filter was transferred to a clean collection tube and 5 μL 0.1 μg/μL trypsin solution together with 95 μL 200 mM ammonium bicarbonate was added to the filter. The filter was incubated overnight in the ThermoMixer with thermotop at 37 °C. Sample collection was performed by centrifuging for 30 min. Subsequently, 150 μL 200 mM ammonium bicarbonate was added to the filter and a centrifugation of 30 min. followed. The samples were again centrifuged for 30 min. after the addition of 150 μL of 10% acetonitrile (ACN), 0.1% formic acid (FA) in MS-H_2_O. Finally, the samples were cleaned and concentrated using solid phase extraction, as described previously. The results from the different sample preparation methods applied to identify extracellular proteins were combined before further analysis for glycosyl hydrolase enzyme candidates. Shotgun proteomics was performed on an EASY 1200 nano-LC coupled to a Q Exactive plus Orbitrap Mass Spectrometer (Thermo Scientific, Germany). Chromatographic separation was carried out on a 0.05 x 150 mm C18 column (Thermo Scientific, catalogue number 164943), with mobile phase A consisting of 0.1% formic acid (FA), 1% acetonitrile (ACN) in MS-H_2_O and mobile phase B consisting of 0.1% FA and 80% ACN in MS-H_2_O. The chromatographic profile included an initial 2-minute run with 5% B, followed by a linear gradient from 5% to 25% B over 90 min, then a gradient to 55% B over 60 min. The gradient was subsequently returned to 5% B within 3 min. and equilibrated for an additional 20 min., all at a constant flow rate of 350 nL/min. Sample injections of 2 μL were performed with blank runs in between each sample. Electrospray ionization operated in the positive mode, and MS1 analysis was conducted at a resolution of 70,000, with an AGC target of 3e6 and a maximum injection time of 75 ms. Applying the data dependent acquisition mode, precursors for fragmentation were isolated using a 2.0 m/z window over a scan range of 385–1250 m/z. A normalized collision energy (NCE) was set at 28%. MS2 spectra were collected at a resolution of 17,500, an AGC target of 2e5 and a maximum injection time of 75 ms. Database searching was performed using PEAKS Studio 10.5 (Bioinformatics Solutions Inc., Canada) using the sequence database obtained from whole metagenome sequencing experiments (described above) and the GPM crap contaminant database. Carbamidomethylation was considered a fixed modification, while oxidation and deamidation were included as variable modifications. Trypsin was set as proteolytic enzyme, allowing a maximum of three missed cleavages. The parent mass error tolerance was set to 20.0 ppm and the fragment mass error to 0.02 Da. Finally, filtering criteria, included a 1% false discovery rate (FDR) for peptide-spectrum matches and a requirement of at least 2 unique peptides for protein identification.

### Functional annotation pipeline

To facilitate the identification of the target glycoside hydrolase, a streamlined bioinformatics pipeline was established. For this, a Virtual Box (Oracle VM Virtual Box, version 7.0.2) was set up with Linux Mint 21 (cinnamon 64 bit) and the pipeline was operated using Python 3.10.12. The functional annotation pipeline was initiated within the python module. The input metagenomics fasta file is first split into smaller files containing max. 1000 proteins each. The program then loops through these files and performs an annotation using HMM sequence models, by using InterProScan (version 5.59-91.0). InterProScan is run from within the python program by making use of the module ‘subprocess’. The split fasta files serve as an input, generating an output in xml format. The precalculated match lookup service of InterProScan has been disabled to run everything locally. After the annotation with InterProScan, a keyword search was performed to mine for sequences that contain elements matching the target functions. Lastly, the data were combined with the metaproteomics results (identified proteins, and abundances).

## Results

### Enrichment on pullulan as the sole carbon source

To enrich for microbes capable of degrading and utilizing pullulan at high temperatures (57°C) and low pH (pH 4.5), pullulan was added as the sole carbon source to the cultures. To evaluate the impact of the inoculum on the enrichment of microbial communities and the enzymes obtained, two enrichment cultures were established. The first, referred to as the ‘compost enrichment’, was inoculated with organic matter from a compost pile. This microbiome was already exposed to elevated temperatures during composting. The second enrichment, called ‘soil enrichment’, was inoculated with soil from the bank of a pond, which was exposed to environmental temperatures (5–20 °C). To eliminate cross contamination between the enrichments, the experiments were conducted under aseptic conditions. Additionally, a control enrichment experiment without any inoculum was included. The compost and ‘soil’ enrichments were maintained for 14 and 20 days, respectively. The cultures were transferred to fresh medium twice during these periods. Daily measurements of OD_600_ and pH were taken, and the pH was adjusted to 4.5 with 0.2 M KOH as needed. After the second transfer, pullulan depletion was monitored daily using high-performance liquid chromatography. Once pullulan was completely depleted (SI document, Figure S1), the enrichments were stopped, and biomass samples were collected for metaproteomic and metagenomic analyses. In pursuit of enzymes capable of functioning at even higher temperatures, one additional enrichment was operated at 75°C. An overview of the enrichment workflow is presented in Figure 1. Two days after inoculation, microscopy confirmed the presence of microorganisms in both the compost and ‘soil’ enrichments, while the control showed, as expected, no signs of growth (Figure 2A). The increase in OD_600_ following both the first and second transfers further indicated successful microbial growth (Figure 2C).The enrichment at higher temperature (75 °C) showed no microbial growth at all (Figure 2A). Over time, the pH of both the compost and ‘soil’ enrichments decreased, necessitating daily adjustments with KOH (Figure 2C). No organic buffer was used to prevent it from becoming a substrate for the microorganisms. Assuming the following biomass growth stoichiometry:

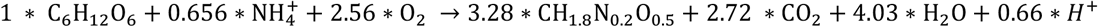

**Figure 1.**
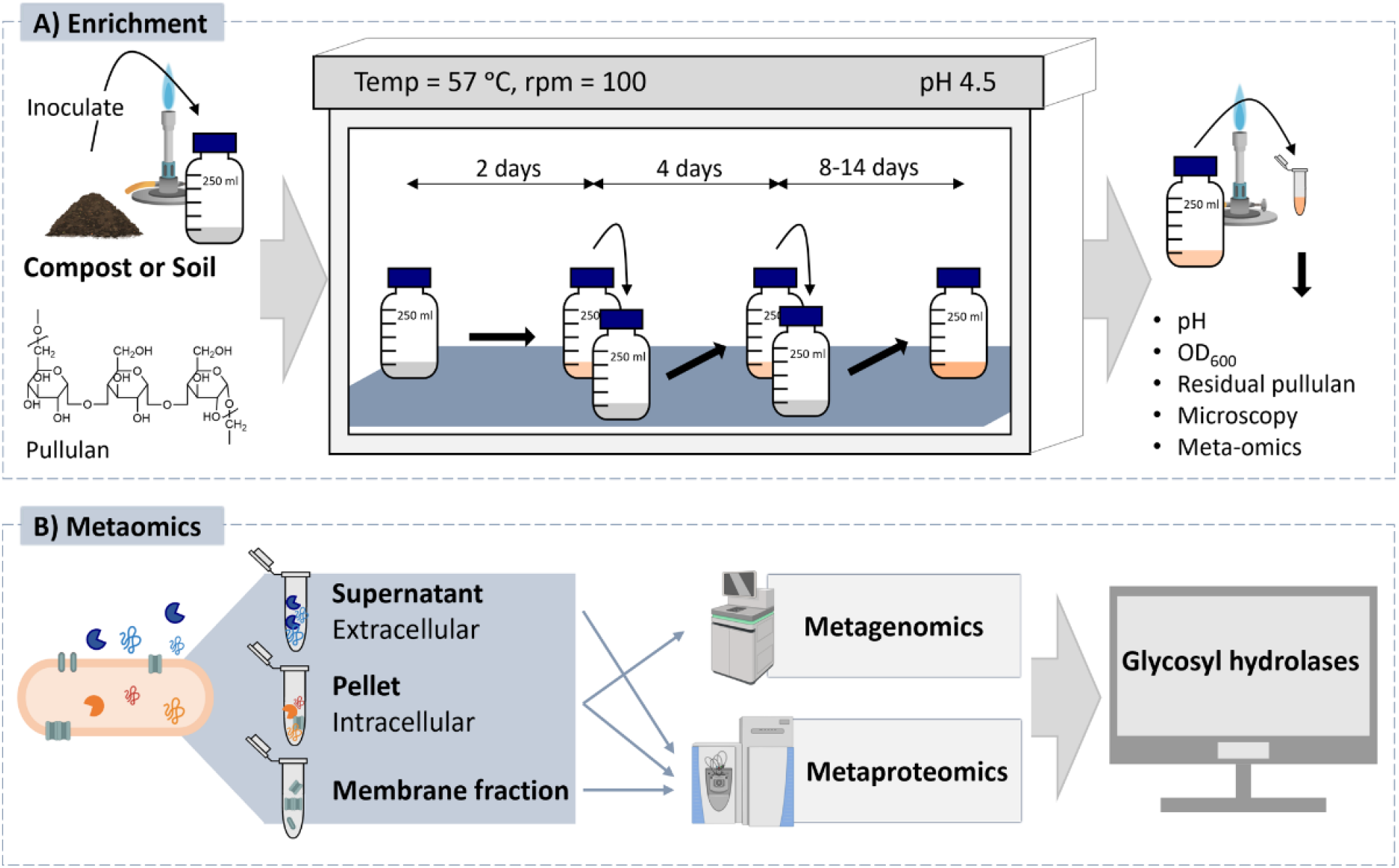
Employed workflow for the discovery of novel glycoside hydrolases from microbial enrichments. **A)** Enrichment cultures at 57 °C and pH 4.5 were established by inoculating either with compost (elevated temperature microbial source) or soil (temperature source control), using pullulan as the sole carbon substrate. In addition, growth medium without inoculum served as a control for cross contamination. The pH, OD_600_ and residual pullulan were monitored daily. The enrichments were studied by microscopy and once all pullulan was depleted the biomass and the secretome were analysed by metagenomics and metaproteomics. **B)** The enrichment biomass was subjected to whole metagenome sequencing. Additionally, metaproteomics analysis was performed on both the cell biomass and supernatant fractions (secretome). While not utilized in this study, future metaproteomic investigations could include a separate analysis of the membrane fraction, offering deeper insights into substrate degradation and update pathways, by identifying dedicated transporters. The subsequent functional annotation pipeline was tailored to identify potential glycoside hydrolase candidates.

**Figure 2.**
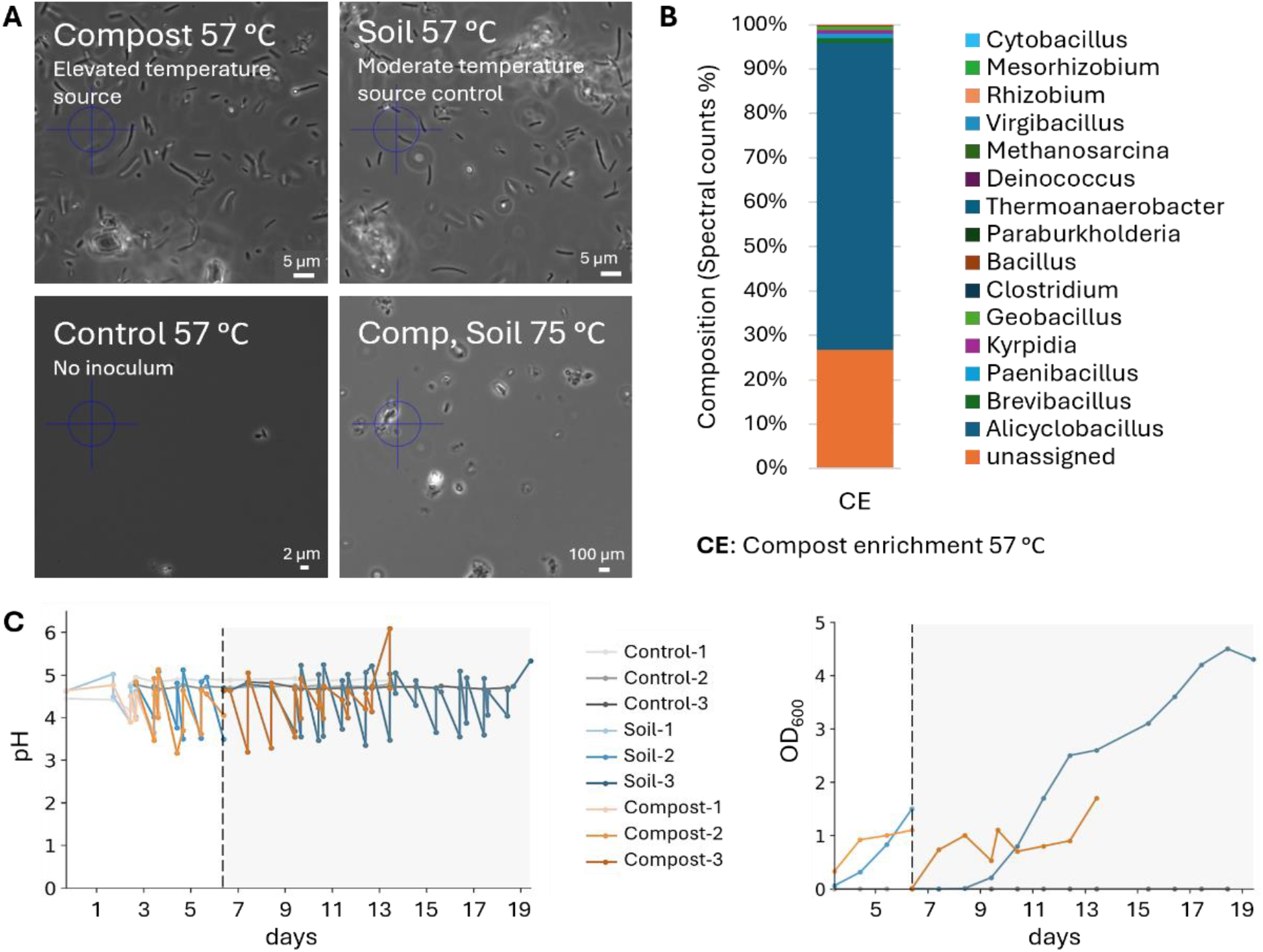
**A)** Microscopy images of the enrichment cultures with a 100x magnification. Compost (elevated temperature microbial source): inoculated with organic matter of a compost pile. Soil (temperature source control): inoculated with sediments from a pond. Control: No inoculum. Comp, Soil 75 °C: inoculated with both compost and soil, incubated at 75 °C. All images were taken two days after inoculation. **B)** Taxonomic composition (genus level) based on spectral counts (metaproteomics) of the ‘Compost’ enrichment biomass, normalized to 100%. Approx. 70% of the counts could be assigned to the genus Alicyclobacillus. The legend shows genera with >10 spectral counts (for full annotation see Figure S4). **C)** pH and OD_600_ measurements during the compost, soil and control enrichment cultures. Grey area: period after the second transfer to fresh medium.

Acidification was likely the result of ammonium uptake (see SI document). Moreover, HPLC-UV analysis using an Aminex organic acid analysis column showed no additional peaks indicative of organic acids, excluding the possibility of acid production during the enrichment (SI document, Figure S1). To verify that pullulan was indeed consumed by the microorganisms, the growth medium was analyzed by HPLC after acid hydrolysis. Specifically, residual pullulan was hydrolyzed to glucose by incubating the medium at 100°C with 4 M Trifluoroacetic acid (TFA). This showed, that over enrichment time, the detected glucose levels decreased in both the compost and ‘soil’ enrichments, whereas the glucose level in the control remained unchanged as expected (SI document, Figure S2).

### *Alicyclobacillus* identified as the dominant genus in both enrichments

The inocula and the biomass from the enrichments were analyzed using whole metagenome sequencing and metaproteomics. Interestingly, the compost inoculum was found to contain a complex spectrum of microorganism based on a taxonomic classification of the metagenomics data. In total, the classification of obtained open reading frames (ORFs) yielded more than 100 different genera, including *Geofilum, Capillibacterium, Methanoculleus*, and *Methanosarcina* (Figure S3).Over 80% of the identified genes could not be classified using this method, pointing to a significant proportion of yet undescribed microbes. This finding highlights the vast diversity and novelty of microorganisms in the inoculum. After enrichment on pullulan at 57°C and pH 4.5, in both enrichments, the complexity reduced to one dominant genus, namely *Alicyclobacillus* (Figure 2B and SI document Figure S4). Thereby indicating that despite the different origin of the inocula, the imposed conditions could select for the same microbe.

### Identification of new glycoside hydrolases from Alicyclobacillus

To identify potential glycoside hydrolases which enable *Alicyclobacillus* to grow efficiently on pullulan, we established a pipeline that integrated functional characterization of the whole metagenome, with the expression and localization data obtained by metaproteomics (Figure 3A). The pipeline first annotates proteins using the InterPro database with a local InterProScan installation, providing an overview of protein families, domains, and sites for each gene. These annotation searched to identify potential enzymes related to hydrolytic activity and pullulan degradation. Keywords included “family 13,” “GH13,” “family 57,” “GH57,” “amylase,” “glycosidase,” “glycoside hydrolase,” and “pullulan”. Unannotated genes were also retained in the final list to allow identification of completely novel sequences. The subsequent integration of metaproteomics data finally provided information on the expression levels and localization of these proteins. While whole metagenomic sequencing of the biomass from the enrichment identified around 14,000 genes (Figure 3B), the use of functional annotations and keyword searches narrowed down the number of potential enzymes to approximately 2,000 genes for the compost enrichment. However, validating the activity of each gene would remain a costly and time-consuming process. Advantageously, integrating metaproteomics data reveals which enzymes are actively expressed. Furthermore, in the case of pullulan, it was also expected that the pullulan degrading enzymes are highly abundant, given their necessity for making the polymer accessible for uptake and growth. Additionally, metaproteomics also provides information on the localization of the proteins. For example, by measuring the biomass pellet and supernatant separately, we could differentiate between intracellular and secreted enzymes. This especially supports the identification of enzymes that are involved in the degradation of the carbon source before cellular uptake^39^. Remarkably, for the compost enrichment, integrating the metaproteomics data reduced the number of enzyme candidates to 96 intracellular and 17 extracellular enzymes, the latter representing potential extracellular pullulan-degrading hydrolytic enzymes (Figure 4). Thus, employing metaproteomics significantly narrowed down the pool of target enzyme candidates. This greatly streamlines subsequent functional and structural investigations, as well as biocatalytic testing after recombinant expression. From the 17 extracellular enzymes, six obtained a functional annotation through the InterPro pipeline, and two of them were identified as containing a domain from the glycoside hydrolase family 13, which includes enzymes such as alpha-amylases and neopullulanases (InterPro IPR006047). Therefore, these sequences are the most likely candidates for possessing a pullulan degrading activity. Nevertheless, some expressed proteins lacked functional annotations from our pipeline and would require also further activity testing.

**Figure 3.**
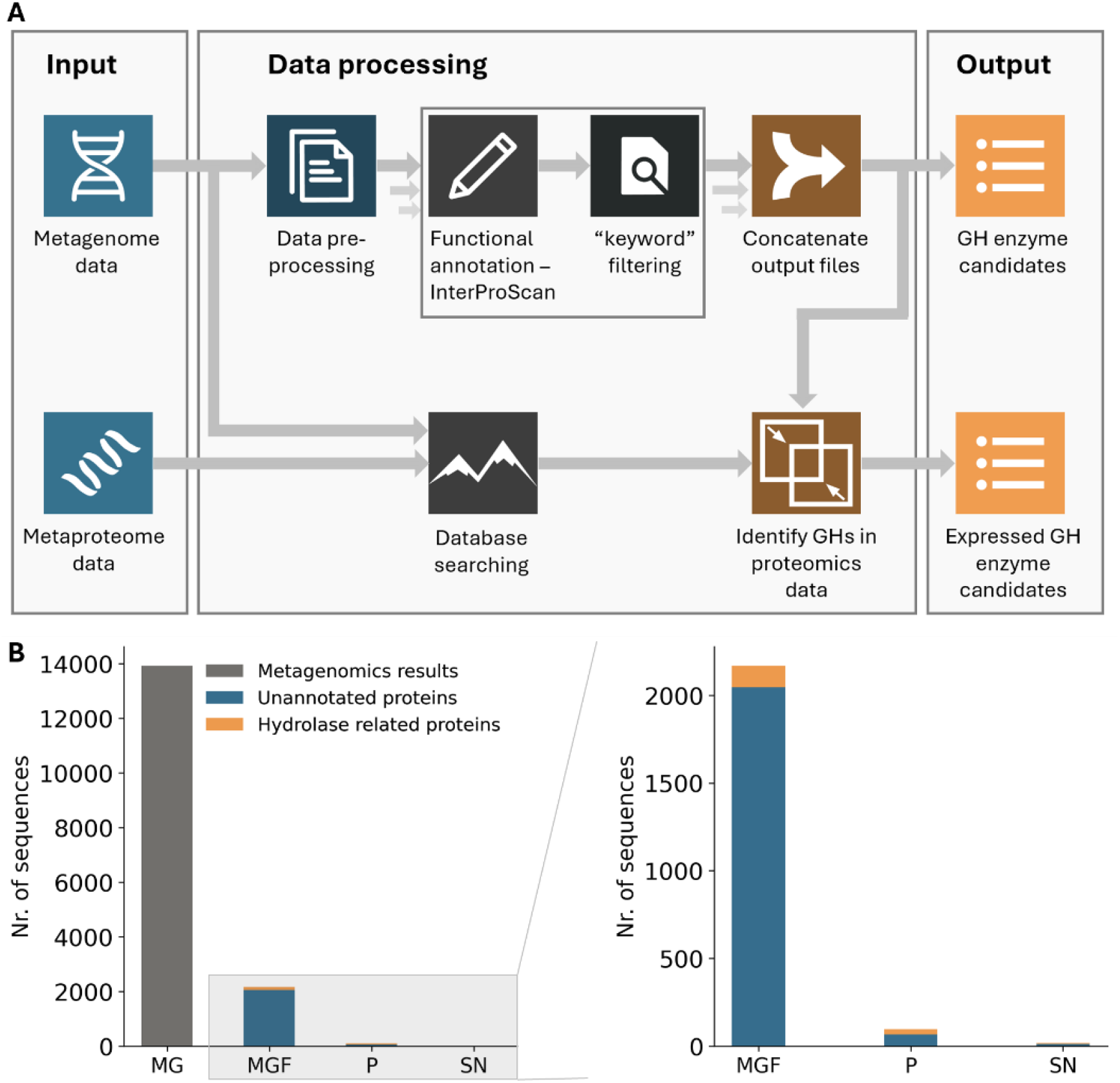
**A)** Schematic overview of the employed bioinformatics pipeline. Functional annotation of the whole metagenome sequencing data was performed using InterProScan. Functional entries containing hydrolase-related keywords (see Table S1) were flagged as potential glycoside hydrolases. The annotated database was subsequently employed to analyse the metaproteomics data by database searching, which ultimately identified a focused number of glycoside hydrolase (**GH**) candidates that were confirmed to be expressed. **B)** The number of pullulan-degrading enzyme candidates obtained from the metaproteomics approach is compared to the total number of sequences obtained from the whole metagenome sequencing data. While metagenomic datasets typically encompass thousands of genes, combining this data with comprehensive functional classification and metaproteomics experiments on both cell pellet and supernatant fractions narrows the focus to a specific set of glycoside hydrolase candidates. In the case of pullulan, this approach also confirmed a two-step degradation process occurring both extracellularly and intracellularly. **MG**: number of genes obtained from the ‘compost’ enrichment by whole metagenome sequencing, **MGF**: number of GH candidates based on comprehensive functional classification of metagenomics data, **P**: number of GH candidates based on metaproteomics on the cell pellet, **SN**: number of GH candidates based on metaproteomics on the supernatant fraction (secreted proteins).

**Figure 4.**
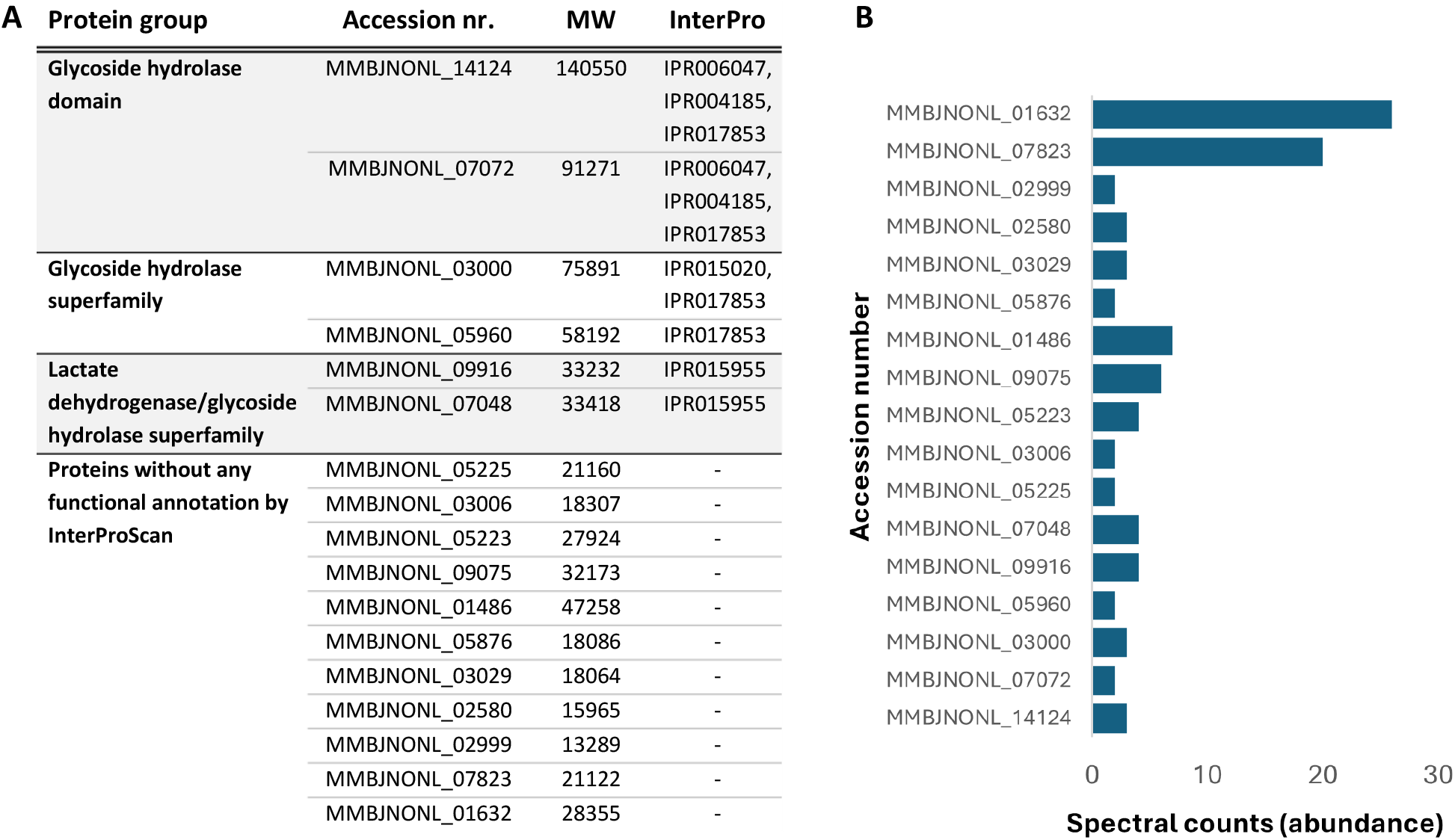
Identified pullulan degrading enzyme candidates in the supernatant (secretome) of the compost enrichment using metaproteomics. **A)** InterProScan annotations. IPR06047: Glycoside hydrolase, family 13, catalytic domain; IPR004185: Glycoside hydrolase, family 13, N-terminal Ig-like domain; IPR017853: Glycoside hydrolase superfamily; IPR015020: Rv2525c-like, glycoside hydrolase-like domain; IPR015955: Lactate dehydrogenase/glycoside hydrolase, family 4, C-terminal (homologous superfamily). **B)** The graph shows the spectral counts based abundance of the identified proteins in the supernatant fraction (secreted proteins).

The dominant species identified in both the compost and ‘soil’ enrichment is *Alicyclobacillus acidocaldarius*, which was first isolated from an acid thermal environment in Yellowstone national park by Darland and Brock^40^. Interestingly, this bacterium has also been reported for causing spoilage in the fruit juice industry^41^. *Alicyclobacillus acidocaldarius* is a spore-forming, rod-shaped bacterium that typically grows at temperatures of 45–70°C (optimum 60–65°C) and pH values of 2–6 (optimum 3–4), closely matching the applied enrichment conditions. Later, Matzke and co-workers identified a gene (CdaA) encoding a cyclomaltodextrinase (neopullulanase)^42^. This cytoplasmic protein was able to hydrolyze the 1,4-glycosidic linkages of pullulan, forming panose as a product. In addition, they identified the gene AmyA encoding an enzyme with amylopullulanase activity, enabling the hydrolysis of the 1,6-glycosidic linkage of pullulan, forming maltotriose. Therefore, AmyA likely hydrolyzes pullulan into maltotriose units extracellularly, which subsequently can be transported over the membrane. CdaA then further hydrolyzes maltotriose to glucose which can enter the central carbon metabolism (Figure 5A). Although Matzke and colleagues already identified two pullulan-degrading enzymes in *Alicyclobacillus*, the elevated temperature and acidic pH in our enrichment yielded a strain with only moderate sequence homology to the previously identified one. Sequence alignment of the AmyA and CdaA genes identified by Matzke and coworkers against the metagenomics data from the biomass of the compost enrichment resulted in six matches (Figure 5B). The best match showed only 80% sequence identity. Furthermore, the matches with high sequence identity to AmyA (MMBJNONL_07072 and MMBJNONL_14124 in Figure 5B) were identified as extracellular by the enzyme discovery approach, while the best match with CdaA (MMBJNONL_12997 in Figure 5B) was classified as intracellular. Interestingly, sequence alignment of the AmyA and CdaA genes against the metagenomic data of the compost inoculum yielded matches with only ∼45% sequence identity, while the enzymes identified through the employed enrichment metaproteomics approach were not detected at all (Figure 5). Although deeper genomic sequencing might detect these enzymes, this underscores the effectiveness of combining microbial enrichment with metaproteomics.

**Figure 5.**
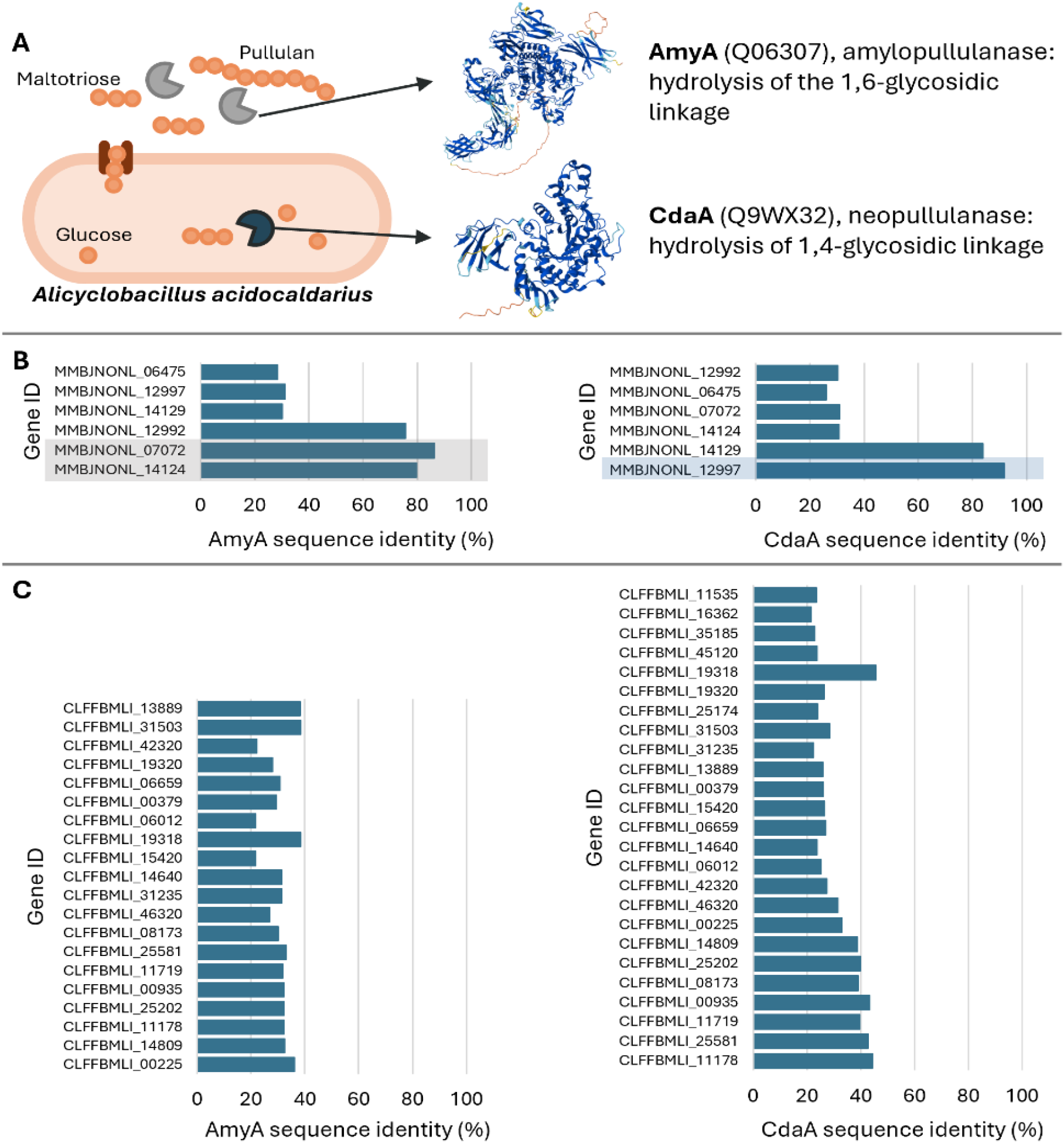
**A)** Proposed pullulan degrading pathway in *Alicyclobacillus* based on the identified enzymes in this study and recent work performed by Matzke et al. (2000). AmyA hydrolyses the 1,6-glycosidic linkage of pullulan extracellularly. The maltotriose units are then imported, after which CdaA hydrolyses the 1,4-glycolytic linkage to form glucose, which can enter the glycolytic pathway. **B)** Sequence alignment of the AmyA (Q06307) and CdaA (Q9WX32) gene, respectively, against the metagenomics data of the compost enrichment culture. Genes showing high sequence identity and which were identified with the glycoside hydrolase bioinformatics pipeline are highlighted in grey (extracellular), or in blue (intracellular). The identified genes with high sequence identity (MMBJNONL _12992, MBJNONL_07072, MMBJNONL_14124, MMBJNONL_ 12997 and MMBJNONL_14129) were not identified in the metagenomics data of the inoculum, which demonstrates the effectiveness of performing microbial enrichment combined with metaproteomics.**C)** Sequence alignment of the AmyA (Q06307) and CdaA (Q9WX32) gene, respectively, against the metagenomics data of the compost inoculum.

## Discussion and conclusions

Advancements in next-generation sequencing have opened up new avenues for the discovery of novel enzymes from microbial sources^28^. However, a purely sequence-based approach is time consuming and commonly overlooks enzymes with new sequences, and the alternative function-based approach is costly and requires the availability of high-throughput activity assays^2^. Therefore, methods that narrow down the pool of potential enzyme candidates without losing the ability to find completely novel enzymes are of great advantage. The combined application of enrichments with multi-omics allows to search for completely novel enzymes operating at defined conditions. We exemplified this approach by searching for new pullulan-degrading enzymes capable of operating at elevated temperatures and acidic pH, from two complex microbial sources (i.e. organic matter from compost, and soil from a pond). The enrichment used pullulan as the sole carbon source, ensuring that only microorganism with pullulan-degrading and utilizing capabilities would thrive. In addition to selecting for a desired function, this process allows customization of conditions to match industrial requirements, such as temperature, pH, and salinity. Nevertheless, the enrichment approach is limited to enzymes that are required for cellular growth or survival, such as enzymes involved in carbon or nitrogen source utilization. Following enrichment, the biomass can be analyzed by whole metagenome sequencing and metaproteomics. Whole metagenome sequencing alone identified approximately 14,000 genes in the compost enrichment, which was narrowed down to around 2,000 candidates after functional classification. Integrating metaproteomics data further allowed to focus on actively expressed enzymes and their specific locations, such as intracellular, membrane-bound, or extracellular. For the compost enrichment, this approach reduced the number of enzyme candidates to 96 intracellular and only 17 secreted enzymes.

Nevertheless, the presence of dead and consequently lysed cells, can result in the identification of intracellular proteins in the supernatant. To assess the amount of lysed cells we compared the proteins identified in the supernatant with the 50 most abundant proteins in the biomass pellet (determined by spectral counts). However, 18 of these proteins were not at all identified in the secretome, and a significant cytoplasmic contamination is unlikely. Also, the supernatant only showed traces of a few of the enzymes of the abundant glycolytic pathway (glyceraldehyde-3-phosphate dehydrogenase, phosphoglycerate mutase and glucose-6-phosphate isomerase), which further confirms that there was no significant cell lysis taking place. Finally, some proteins (e.g. membrane proteins) are difficult to detect by conventional proteomics approaches, and might be missed without additional optimisation of the sample preparation^43^. Despite *Alicyclobacillus* being identified as the dominant genus, other coexisting organisms, though present at lower abundances, were also identified. Therefore, it cannot be excluded that also these organisms are involved in hydrolyzing pullulan, which subsequently can be consumed by multiple microbes. Intriguingly, a third enrichment was initiated at 75 °C, but no growth was observed. A review by Kahar et al. highlights diverse pullulan-degrading enzymes identified from microbial sources, demonstrating activity at high temperature and low pH^13^. For instance, pullulanases from organisms like *Fervidobacterium nodosum* Rt17-B1 (optimal temperature, T_opt_ of 80°C and an optimal pH, pH_opt_ of 5), *Desulfurococcus mucosus* DSM 2162T (T_opt_ = 85 °C, pH_opt_= 5), and *Thermoanaerobacter* sp. B6A (T_opt_ = 75 °C, pH_opt_ = 5) are some examples. However, enzyme activity at 75 °C and pH 4.5 doesn’t guarantee that the organism can grow under these conditions. For instance, for *Fervidobacterium nodosum* the growth range is restricted to pH 6.0–8.0. Moreover, it is an obligate anaerobe, preventing growth under the conditions set in this study^44^. Similarly, *Desulfurococcus mucosus* and *Thermoanaerobacter* are both obligate anaerobes and do not grow under aerobic conditions^45,46^. One enrichment was inoculated with compost (“compost” enrichment), which typically has an elevated temperature and is therefore expected to harbor microorganisms that can thrive under such conditions. To assess whether these naturally high temperatures are essential for enriching microbes which thrive at elevated temperatures, a second enrichment was inoculated using soil from a pond (‘soil’ enrichment). While the inocula displayed a complex and diverse microbial composition, *Alicyclobacillus* emerged as the dominant genus from both inocula. Cross-contamination is unlikely, as no growth was observed in the control experiment. These findings suggest that despite different initial temperatures of the inocula, similar organisms thrived under identical enrichment conditions. The similarity may also be attributed to the proximity of the sampling sites, approximately 20 kilometres apart, and their shared soil environment. Both pullulanase types I and II often have the ability to degrade starch, a common natural plant polymer found in soil, which might support the growth of similar organisms. Although previous studies identified two pullulan-degrading enzymes, AmyA and CdaA, in *Alicyclobacillus*, our enrichment metaproteomics approach revealed distinct enzyme sequences with potentially altered properties (Figure 5). These identified enzymes were not detected in the whole metagenome sequencing data of the inoculum, highlighting the effectiveness of our approach. Metaproteomics confirmed that the putative AmyA is extracellular, while the putative CdaA is intracellular. This suggests that AmyA hydrolyzes pullulan into maltotriose units extracellularly, which are then imported into the cell and further broken down by CdaA into glucose. Supporting this, we also found a maltose/maltodextrin-binding periplasmic protein and the maltose/maltodextrin transport system MalG being expressed. This aligns with previous findings of a high-affinity, binding-protein-dependent ABC transport system specific for maltose and maltodextrins in *Alicyclobacillus*^*47*^.

In summary, we demonstrated the combination of microbial enrichment and metaproteomics for discovering novel microbial glycoside hydrolases, as exemplified by pullulan enrichment from two different microbial sources. The integration of metaproteomics significantly accelerated enzyme identification in these complex environments, reducing the number of genes requiring subsequent biochemical validation.

## Supporting information

SI DOC

SI EXCEL DOC

## Author contributions

JvE, MvL, GvK and MP conceived and designed the project. JvE and SvS performed the experiments. GvK and MP provided critical input to experiments and/or protocols. JvE, SvS and MP conducted the data analysis. JvE and MP generated the figures and wrote the original draft. All coauthors contributed to reviewing and editing the manuscript.

## Acknowledgements

The authors acknowledge Dita Heikens for her support with sample preparation and all other colleagues from the Department of Biotechnology for valuable discussions. This work was supported by the TU Delft Department of Biotechnology Zero Emission Biotechnology Program (ZEB). The authors acknowledge IFF for providing pullulan and covering the Illumina short-read sequencing of the enrichment.

