## Supplementary material for "Discovery of microbial glycoside hydrolases via enrichment and metaproteomics": SI DOC

### S1. Culture acidification

Over time, the pH of both the 'compost' and 'soil' enrichments decreased, necessitating daily adjustments with KOH (Figure 2C). This acidification was likely a result of ammonium uptake for biomass growth. Initially, each enrichment contained 0.5 g of pullulan (which equals 3.09 mmol of glucose). Throughout the enrichment process, approximately 2.61 ml of 0.2 M KOH (0.52 mmol KOH) was added to maintain the pH at 4.5. Thus, for every mole of glucose consumed, 0.17 mol of KOH was needed to neutralize the produced $H^+$, indicating that 0.17 mol of $H^+$ was generated during growth. The following biomass growth stoichiometry (normalized to the consumption of 1 mol glucose) is assumed:

$$1 * \mathrm{C_6H_{12}O_6} + 0.656 * \mathrm{NH_4^+} + 2.56 * \mathrm{O_2} \rightarrow 3.28 * \mathrm{CH_{1.8}N_{0.2}O_{0.5}} + 2.72 * \mathrm{CO_2} + 4.03 * \mathrm{H_2O} + 0.66 * H^+$$

According to this equation, the consumption of 1 mol of glucose could produce up to 0.66 mol of $H^+$. The observed acidification in the culture falls within the theoretically possible range of acidification.

### S2. HPLC chromatograms showing the Pullulan depletion

To confirm that the microorganisms were growing on pullulan, the pullulan depletion was monitored over time by HPLC, both before (**Figure S1**) and after complete hydrolysis (**Figure S2**).

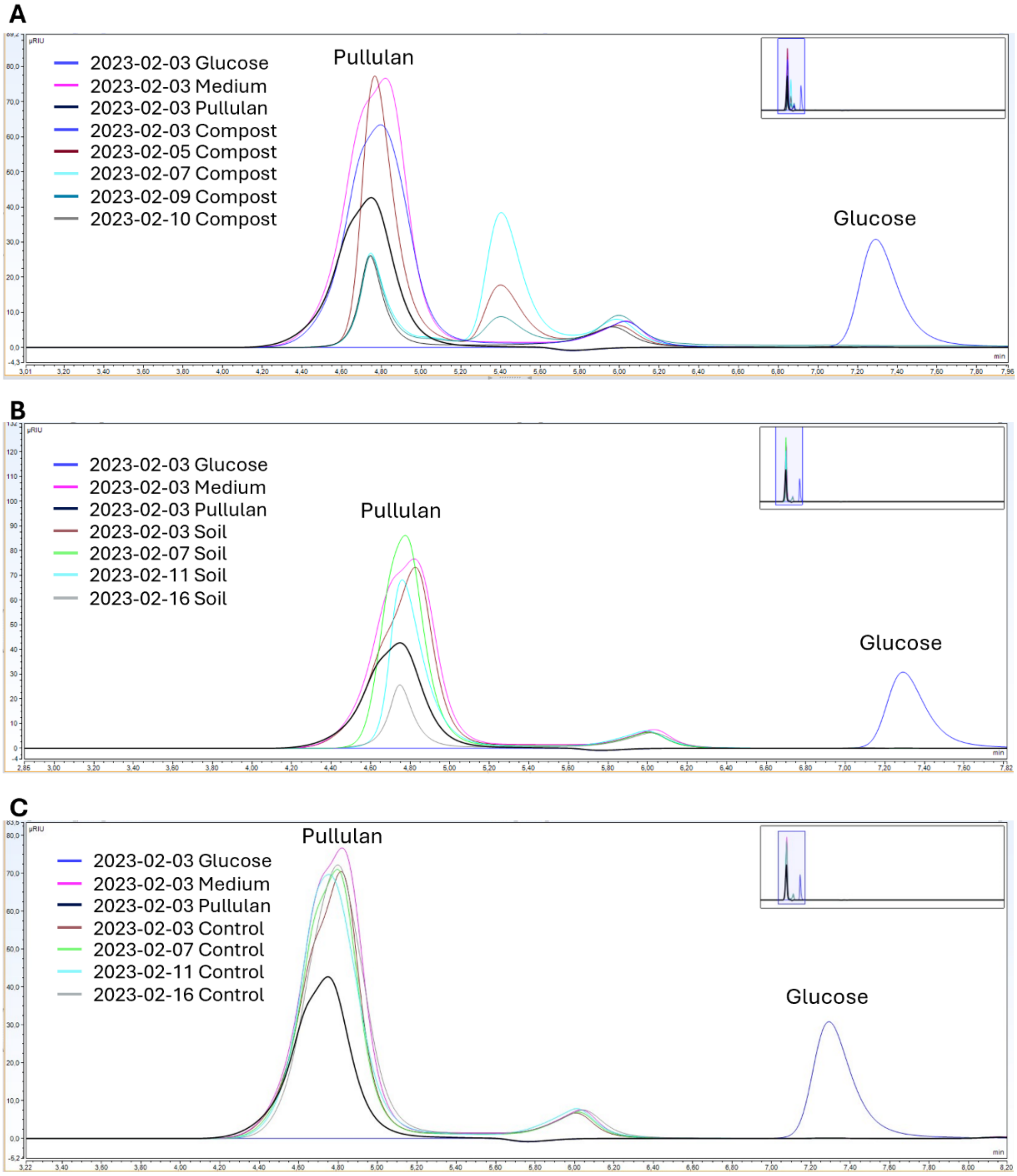

***Figure S1.*** Pullulan depletion. Overlayed HPLC chromatograms of the analysed enrichment cultures to follow the pullulan depletion over time. Whole broth samples were centrifuged after which the supernatant was analysed. HPLC conditions are described in the method section. **A)** Compost enrichment (elevated temperature microbial source): inoculated with organic matter of a compost pile. **B)** Soil enrichment (temperature source control): inoculated with sediments from a cold water pond. **C)** Control: No inoculum.

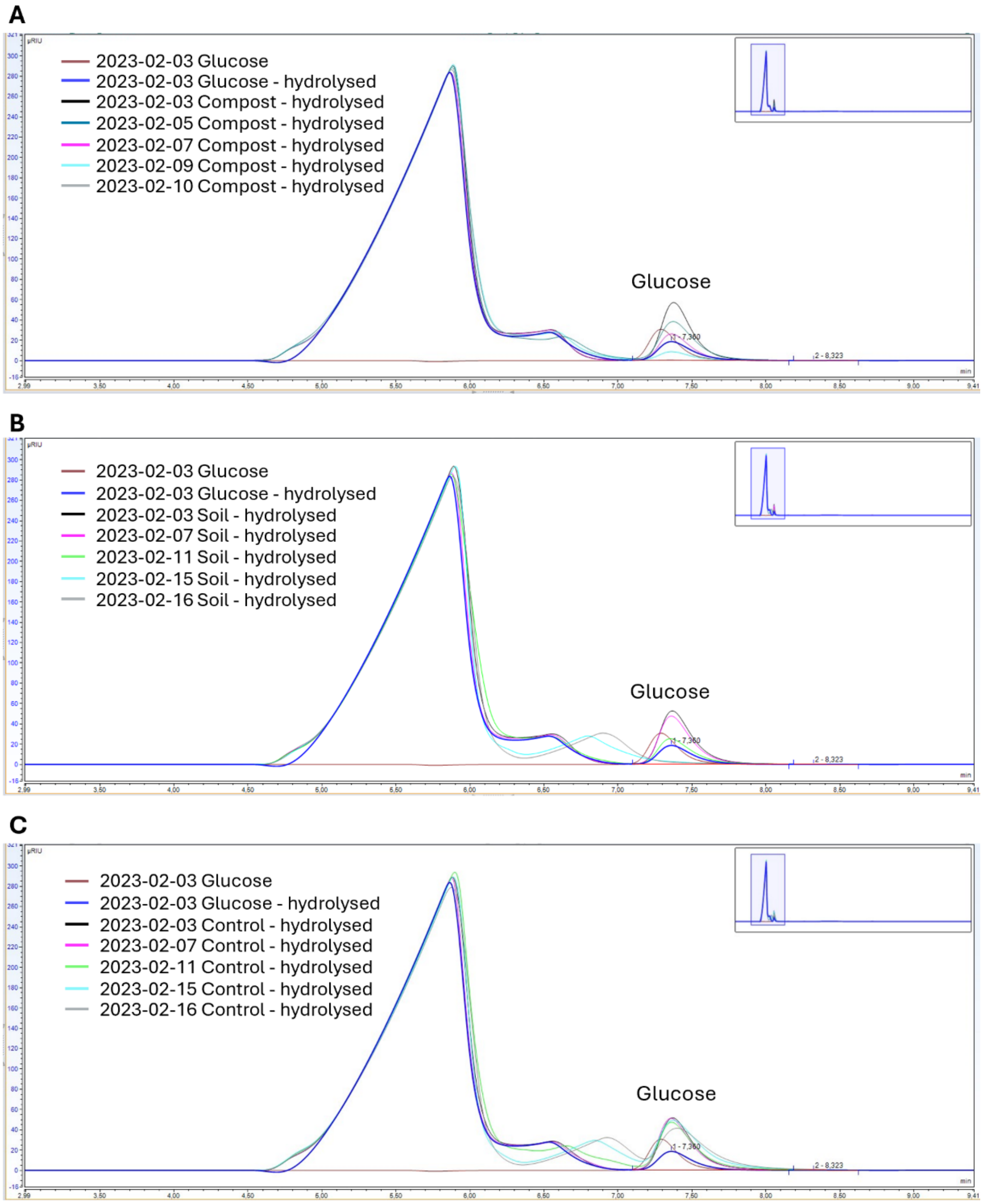

***Figure S2.*** Glucose depletion. Overlayed HPLC chromatograms of the analysed enrichment cultures to follow the glucose depletion over time. Whole broth samples were centrifuged, after which the remaining pullulan in the supernatant was hydrolysed by adding trifluoroacetic acid (TFA) to a final concentration of 4M. Samples were incubated at 100 °C for 4 hours. HPLC conditions are described in the method section. **A)** Compost enrichment (elevated temperature microbial source): inoculated with organic matter of a compost pile. **B)** Soil enrichment (temperature source control): inoculated with sediments from a cold water pond. **C)** Control: No inoculum.

### S3. Taxonomic composition of the enrichment cultures and the inoculum

A taxonomic classification of the compost inoculum (Figure S3) based on the metagenomics data has been performed. For illustration purposes, only the genera with ≥ 5 spectral counts are depicted. The total amount of genera identified by taxonomic profiling equals 421, emphasizing the enormous variety of microorganisms present. A significant number of sequences remains unassigned. Therefore the taxonomic distribution might not be the best representation of the reality and needs to be further optimized. Nevertheless, it does show the enormous variety of microorganisms present in the inoculum. In addition, a taxonomic classification of the enrichment cultures was performed based on the metaproteomics data (Figure S4).

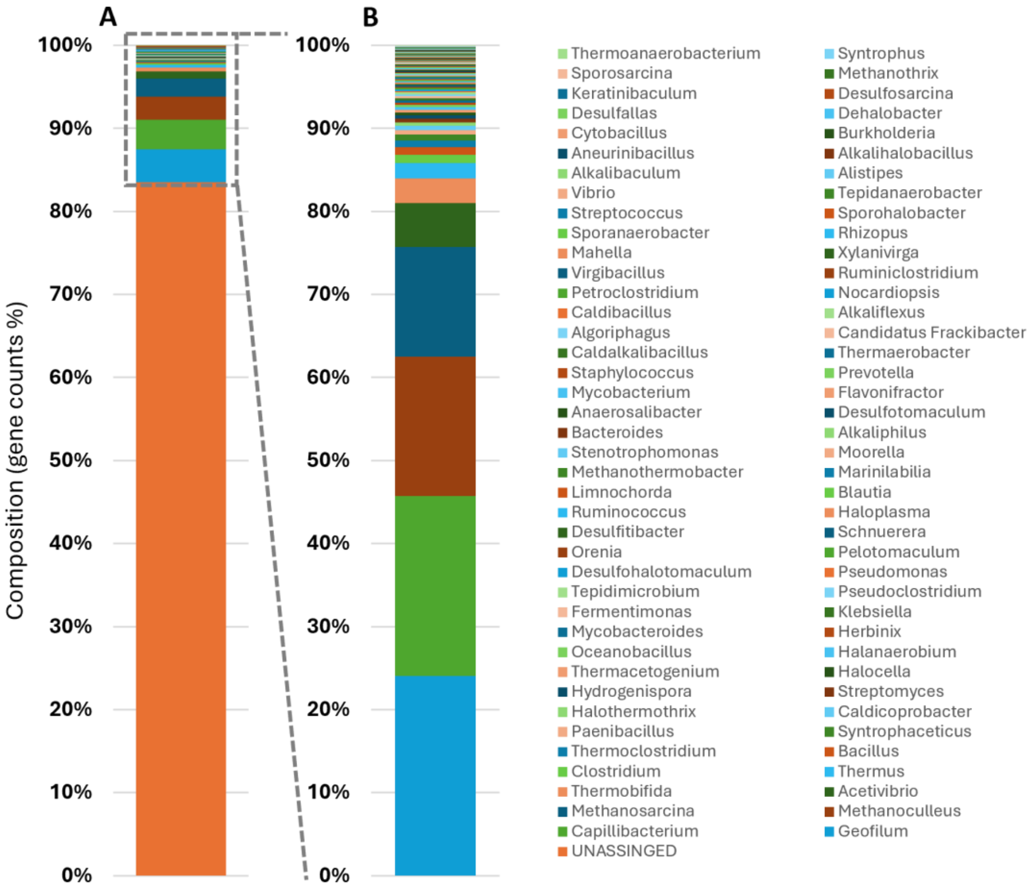

***Figure S3.*** Taxonomic composition of the compost inoculum based on gene count (metagenomics data). Taxonomic classification was performed on gene level. **A)** Including the unassigned sequences; **B)** Excluding the unassigned sequences.

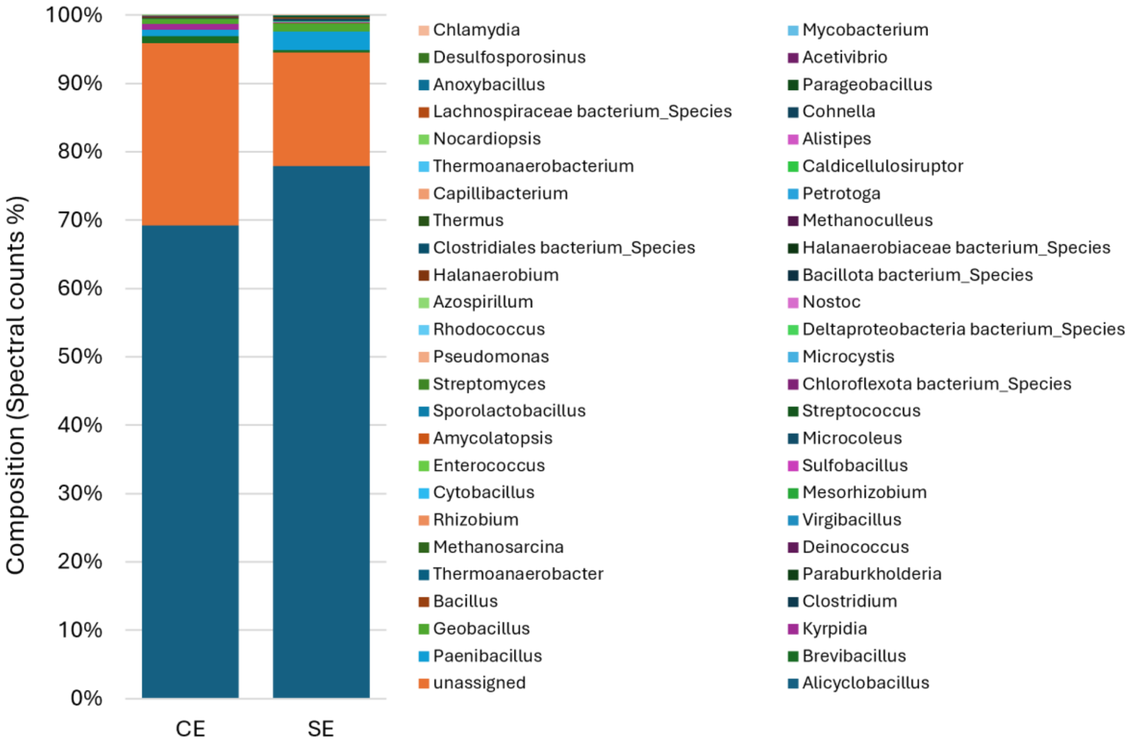

***Figure S4***. Taxonomic composition (genus level) of the 'Compost' and 'Soil' enrichments based on spectral counts (metaproteomics), normalized to 100%. **CE:** Compost enrichment; **SE:** soil enrichment.
